# Estimating the Inter- and Intra-Rater Reliability for NASH Fibrosis Staging in the Presence of Bridge Ordinal Ratings with Hierarchical Bridge Category Models

**DOI:** 10.1101/2021.10.27.466144

**Authors:** Joshua Levy, Carly Bobak, Nasim Azizgolshani, Xiaoying Liu, Bing Ren, Mikhail Lisovsky, Arief Suriawinata, Brock Christensen, James O’Malley, Louis Vaickus

## Abstract

The public health burden of non-alcoholic steatohepatitis (NASH), a liver condition characterized by excessive lipid accumulation and subsequent tissue inflammation and fibrosis, has burgeoned with the spread of western lifestyle habits. Progression of fibrosis into cirrhosis is assessed using histological staging scales (e.g., NASH Clinical Research Network (NASH CRN)). These scales are used to monitor disease progression as well as to evaluate the effectiveness of therapies. However, clinical drug trials for NASH are typically underpowered due to lower than expected inter-/intra-rater reliability, which impacts measurements at screening, baseline, and endpoint. Bridge ratings represent a phenomenon where pathologists assign two adjacent stages simultaneously during assessment and may further complicate these analyses when *ad hoc* procedures are applied. Statistical techniques, dubbed *Bridge Category Models*, have been developed to account for bridge ratings, but not for the scenario where multiple pathologists assess biopsies across time points. Here, we develop hierarchical Bayesian extensions for these statistical methods to account for repeat observations and use these methods to assess the impact of bridge ratings on the inter-/intra-rater reliability of the NASH CRN staging scale. We also report on how pathologists may differ in their assignment of bridge ratings to highlight different staging practices. Our findings suggest that *Bridge Category Models* can capture additional fibrosis staging heterogeneity with greater precision, which translates to potentially higher reliability estimates in contrast to the information lost through *ad hoc* approaches.

## Introduction

Nonalcoholic steatohepatitis (NASH) is a fatty liver condition hallmarked by excessive lipid accumulation. Lipid accumulation can lead to inflammation, hepatocyte ballooning, subsequent necrosis and stellate cell activation, spurring the production of collagen (fibrosis) and leading to progressive, potentially irreversible liver damage. Progression to cirrhosis (end stage fibrosis) can lead to liver failure, the need for liver transplant or death. Advanced fibrosis / cirrhosis also markedly increases the chance of developing hepatocellular carcinoma (HCC)^1–5^ NASH is correlated to western diet, obesity and metabolic syndrome and is set to become the most common cause of cirrhosis, eclipsing viral hepatitis ^6^. However, despite intense study, causal genetic links have not been established, despite associative findings ^7^. Current drugs used to reverse fibrosis progression largely target fibrotic pathways, though none have completed stage 2 clinical trials and gained FDA approval ^8^.

The deficiency in completed NASH clinical trials may be partially attributed to the need for a histopathological clinical endpoint for assessing fibrosis regression / progression from liver biopsy. Accurate fibrosis staging is important for establishing clinical screening criteria to identify which patients may benefit from treatment, as well as determining fibrosis baseline and endpoints. Clinical tests assume that fibrosis endpoints are measured accurately, leaving little room for uncertainty and ambiguity, both within and between raters. However, when variability in ratings exists between pathologist raters, this may signify the presence of several forms of measurement uncertainty, including measurement error, which may reduce study statistical power. These reductions in power significantly hinder efforts to evaluate the true efficacy of NASH treatments. For example, numerous studies have highlighted the high inter- and intra-rater reliability of the NASH CRN staging scale (e.g. carefully chosen representative tissue samples and scoring performed by experts with decades of experience as opposed to variable tissue quality interpreted by a generalist)^9–16^. It is feasible that the described qualitative scoring system, which was designed for clinical prognostication, is not reproducible or reliable enough to rigorously gauge fibrosis regression in a drug trial.

Here, we report on the impact of an additional dimension of uncertainty in the NASH fibrosis scoring system: bridge categories. Bridge ratings occur when the pathologist is unable to determine which of two adjacent stages to assign (e.g., 2 and 3), thereby leading them to report both (e.g., 2-3). This may occur for many reasons: the pathologist has identified features truly indicative of both adjacent stages, the pathologist feels they have insufficient tissue or wants to hedge against the possibility of a more advanced stage, or due to the spatial variability in suggestive prognostic features across the slide. While there are many reasons why a bridge rating may occur, these ratings represent a potential source of variation in the NASH fibrosis scoring system and other such ordinal rating systems ^17^.

Bridge ratings are rarely reported in the literature yet occur frequently in studies that report such ratings. For instance, disease staging systems analogous to the NASH system (e.g., TNM staging ^18,19^) actively discourage pathologists from reporting bridge ratings in their clinical staging manuals without substantiating the reasons for making these *ad hoc* adjustments. *Ad hoc* adjustments include assigning the lower of two stages (downstaging), the higher of the two stages (upstaging), randomly assigning stage (random staging), removing the case altogether from the study or reporting the stage assigned by a consensus of pathologists. Bridge ratings have complicated studies which use the NASH CRN scale for the assessment of the final outcomes, and when reported, researchers have previously resorted to these *ad hoc* procedures^20^.

Performing *ad hoc* adjustments and failing to report bridge ratings may lead to significant bias and/or underestimation of the reported effect for a clinical study. Moreover, these ad hoc score adjustments ignore the fact that bridge ratings present meaningful information which may better assess the efficacy of a drug when the reasons a bridge rating was used are understood. Alternatively, when the mechanism for bridge rating assignment is misinterpreted or bridge ratings are omitted altogether, information is lost.

Recently, new statistical modeling procedures have been developed (referred to as the *Bridge Category Models*) which are able to account for bridge ratings under the different scenarios in which they arise ^17^. However, there do not yet exist methods which can account for multiple raters and repeat observations. Repeat observations represent a statistical phenomenon where, for instance, a group of pathologists may stage a single specimen or shared cohort of specimens. In this very common arrangement, ratings on the same case from different pathologists may differ based on their prior experience, workload, education, etc. Additionally, repeated observations may occur when the same pathologist rates the same case twice and may disagree with themselves ^21–23^ These repeated measurements are accounted for in statistical models which can calculate the inter- and intra-rater reliability for measurement scale validation. Nonetheless, there is no standard procedure for accounting for bridge ratings when reporting inter- and intra-rater reliability.

The aims of this study are thus twofold. First, we present an extension to the *Bridge Category Model* which is able to capture NASH fibrosis stages from multiple raters across multiple timepoints. We use the *Bridge Category Models* to report on the reliability of NASH fibrosis stages between pathologist raters and across time. We compare the results of the bridge category models with that of the *ad hoc* procedures to understand their impact on estimating the variation in ratings. Finally, we compare the observed variability with that reported in the literature and assess how pathologists may differ in their practices for assignment of bridge ratings.

## Materials and Methods

### Data Collection

A total of 287 liver biopsy tissue specimens (wedge resection, fine needle, and core needle) were collected from a cohort of 273 individuals with a proven NASH diagnosis from Dartmouth Hitchcock Medical Center. FFPE tissue blocks were obtained and 5-micron thick sections were cut. Adjacent sections of tissue were stained with hematoxylin and eosin and separately with trichrome stain for fibrosis assessment. Biopsies were scanned using the Leica Aperio-AT2 (Leica Microsystems, Buffalo Grove, IL) scanner at ×20 magnification and stored in SVS image format (JPEG compression at 70% quality).

### Rating Cases

Images of the Trichrome stain from all tissue sections were presented to four pathologists, who independently examined the tissue for fibrosis stage on two separate occasions with an appropriate washout period of at least two weeks, with randomization of study number and image order.

### Modeling Procedures

We utilized two-way mixed effects ordered probit models to assess the reliability of stage assignments both across pathologists (inter-rater) and within pathologist, across two timepoints (intra-rater) ^24–29^. We developed and applied hierarchical *Expanded* and *Mixture* Bridge Category Models, an extension of previously published work, where bridge ratings account for the following scenarios: 1) assignment of an intermediate stage between two existing adjacent stages may better capture the disease pathology (*Expanded*) and 2) ratings from a true stage are blurred across two adjacent stages to hedge against the possibility of incorrect assignment (*Mixture*). We compared results to that obtained through *ad hoc* procedures (down-staging, up-staging, random assignment of up/down-staging for the adjacent stages). The statistical models estimate three variance components after sampling of the posterior distribution of parameters using Hamiltonian Monte Carlo (HMC) procedures ^30,31^:

1. The variance component attributed to different biopsies: 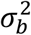. Biopsies are nested within patients and patients are drawn from a large population of patients of interest.
2. The variance component attributed to different pathologists: 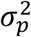. The four pathologists in our study are assumed to have been selected from a population of pathologists that could have staged the biopsies.
3. The variance component attributed to different factors of the interaction between pathologists and biopsies: 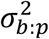. Biopsies are cross-classified with pathologists since every pathologist stages every biopsy ^32^. However, some biopsies may be easier to stage than others in the sense that pathologists yield more similar ratings for them compared to other biopsies. We used the intraclass correlation coefficient (ICC) to communicate inter- and intra-rater measurement reliability, to be detailed in the following sections. The ICC, related to the weighted Kappa statistic (asymptotically equivalent) used in Fibrosis staging studies ^33^, is a score between 0 (no agreement) and 1 (complete agreement) which represents the amount of agreement between (inter)/within (intra) pathologists in their staging given a certain number of replicated measurements. ICC values below 0.75 are considered to be poor to moderate, while estimates above 0.75 are considered to be good. Scores above 0.9 are said to demonstrate clinical validity ^34^. We adopted methods for concurrent estimation of both inter- and intra-rater reliability to communicate our study findings ^35^. To estimate the variance components that explain staging patterns for estimation of the ICC, ordinal regression models estimate the probability of observing a certain stage given the data while taking into account the ordered nature of the stage outcomes. When bridge categories do not exist, the probability of observing a stage *j* is given by: *P(X)*. When bridge categories exist, we must account for the fact that the bridge rating could represent two possible scenarios under which {*j,j* + 1} could be assigned: either the pathologist felt that the lower of the two adjacent stages, *Y = j*, was true but held some belief that the higher rating might be the truth or the pathologist felt that the higher of the two adjacent stages, *Y* = *j* + 1, was true but held some belief that the lower rating might be the truth. Letting *p* represent the probability of the former, the likelihood of the observation {*j,j* + 1} is given by *p *P(X) + (1 −p) *P(X)*. The probability *p* provides meaningful information as to how often the pathologists produce bridge ratings, under the scenario where the pathologist believes the features of the biopsy most likely indicate a lower stage but is hedging against a more serious prognosis ^17^. There exist other specifications of bridge category modeling procedures, though this procedure is of primary interest since interviews with pathologists indicated the greatest concordance with this data generating mechanism. In addition to the aforementioned variance components, the *Mixture* Bridge Category model estimates the following parameter:
4. The probability by which a pathologist assigned a bridge rating under the belief that the lower stage of two adjacent stages was the truth: *p*. When the higher stage is the truth, pathologists assign bridge ratings with a probability 1 − *p*. Pathologists may disagree with each other or with their own prior bridge ratings, so we can report *p* on a rater/test specific basis.

Further details of the hierarchical Bayesian Bridge Category model specification and inference targets can be found in the Appendix (sections “Additional Description of Bayesian Modeling Procedures”, “Complete Bayesian Specification of Hierarchical Mixture Bridge Category Model”, “Description of Priors”). The formulation and derivation of the original *Bridge Category Models* from which this work is based on may be found in a previous publication^17^.

### Assessment of Inter-Rater Reliability

The inter-rater ICC assesses the reliability of stage assignments across pathologists and is given by the following expression:

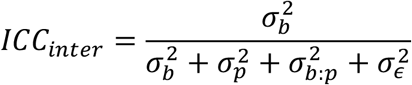

Where 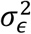 is the individual error variance of the underlying continuous ratings, which are assumed to be normally distribution with a variance of 1 to be consistent with an ordinal probit regression model being used to analyze the categorical rating data - this is one of several ad hoc and imperfect ways of dealing with the lack of an explicit individual error term ^35–37^.

### Assessment of Intra-Rater Reliability

In this study, the intra-rater ICC is assessed by including measurements from two timepoints and capturing the remaining variation that is neither attributed to biopsies nor pathologists. The remaining variation may be attributed to variation within pathologists on subsequent time points. Thus, the intra-rater ICC is given by:

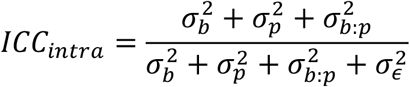

Where here 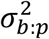 is the variance attributed to the interaction between the biopsy and pathologist ^35^.

### Estimation of Pathologist Bridge Rating Preference where the *Mixture* Model is True

Under the assumptions of the *Mixture* Bridge Category model, rater-specific effects can be reported for proportion *p* for each test/inspection:

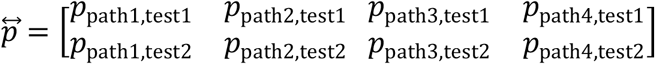

When a bridge rating is assigned, *p*_*pathologist*,*test*_ indicates the proportion of times a specific pathologist assigned two higher stages when they believed the lower stage of the two adjacent stages was true.

### Details of Methods Comparison

In this study, we compare ICC estimates acquired using *Bridge Category Models* to that of *ad-hoc* procedures via reports of 90% equal-tailed credible intervals and perform hypothesis testing with the probability of direction (*pd*), which measures how much of the probability mass of a parameter’s posterior distribution exceeds a target value, where *1 −pd* is analogous to the frequentist p-value ^38^. For the *Mixture Bridge Category Model*, we comment on rater and test-specific estimates of *p*. Specifically, we assign statistical significance of pathologist/test-specific estimates of *p*using the probability of direction and compare the rater/test-specific proportion *p*to 0.5, the nominal bridge rating proportion where no preference exists for up/down staging:

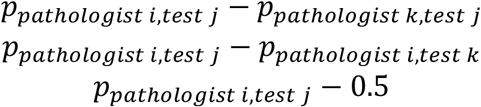

### Software Implementation and Complete Model Specification

The hierarchical models used to estimate the inter- and intra-rater ICC were fit using Bayesian procedures via the Stan programming language, implemented in R via the *Rstan* package ^21,31^. We have made the software for estimating the models available at the following URL: https://github.com/jlevy44/BridgeCategoryNASHFibrosisICC. As noted earlier, a complete Bayesian specification of the modeling procedures and selection of priors may be found in the Appendix.

## Results

### Assignment of Bridge Category Assessments

Of the 2,291 stages assigned by the practicing pathologists, 484 observations (21.1%), were given bridge measurements. Of the bridge measurements, 13.9% of them were stage 0-1, 14.0% were stage 1-2, 45.9% of them were stage 2-3 and 26.2% of them were stage 3-4. Pathologist 1 assigned the lowest number of bridge ratings, (0 during the second test), while Pathologist 3 assigned the most bridge ratings (91/287 or 31.7% of ratings during their second test). Of the 287 biopsies, 230 (80.1%), were assigned at least one bridge rating, which corresponded to 218 out of 273 patients (79.9%, Table 1).

**Table 1:** Breakdown of Bridge Rating Assignments, tabulated over pathologist and test

| Test 1 | Pathologist | Regular Stage Ratings |  |  |  |  |  | Bridge Ratings |  |  |  |  |
| --- | --- | --- | --- | --- | --- | --- | --- | --- | --- | --- | --- | --- |
|  |  | 0 | 1 | 2 | 3 | 4 | Total | 0-1 | 1-2 | 2-3 | 3-4 | Total |
|  | 1 | 19 | 40 | 77 | 55 | 67 | 258 | 0 | 0 | 14 | 15 | 29 |
|  | 2 | 13 | 36 | 78 | 48 | 28 | 203 | 16 | 12 | 38 | 18 | 84 |
|  | 3 | 1 | 13 | 98 | 61 | 31 | 204 | 8 | 12 | 44 | 19 | 83 |
|  | 4 | 65 | 17 | 41 | 36 | 58 | 217 | 15 | 12 | 27 | 14 | 68 |
|  | Total | 98 | 106 | 294 | 200 | 184 | 882 | 39 | 36 | 123 | 66 | 264 |
| Test 2 | 1 | 16 | 43 | 92 | 68 | 68 | 287 | 0 | 0 | 0 | 0 | 0 |
|  | 2 | 66 | 16 | 20 | 41 | 79 | 222 | 7 | 10 | 29 | 16 | 62 |
|  | 3 | 11 | 39 | 56 | 60 | 30 | 196 | 13 | 12 | 40 | 26 | 91 |
|  | 4 | 3 | 18 | 97 | 66 | 36 | 220 | 8 | 10 | 30 | 19 | 67 |
|  | Total | 96 | 116 | 265 | 235 | 213 | 925 | 28 | 32 | 99 | 61 | 220 |
| Combined | Total | 194 | 222 | 559 | 435 | 397 | 1807 | 67 | 68 | 222 | 127 | 484 |

### Inter-Rater Reliability

The inter-rater ICC for fibrosis staging for the *Mixture* model was around 0.754 (CI: 0.668-0.802). The probability that the ICC estimate for the *Mixture* model surpassed other *ad hoc* ICC estimates and was as high as 82.3%, while ICC estimates given by the *Expanded* model were more similar to the *ad hoc* procedures. For instance, the random staging ICC estimate was found to be approximately 10% lower than using the *Mixture* approach (Table 2). The posterior intervals of ICCs for the *Bridge Category Model* approaches (Mixture, Expanded) were significantly lower than the *ad hoc* procedures, indicating greater precision.

**Table 2:**
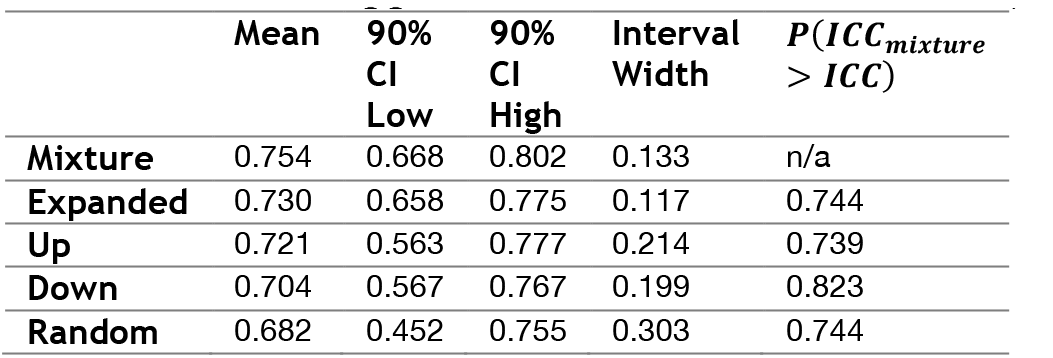
Inter-Rater ICC estimates for all statistical approaches and probability of how much ICC for *Mixture* approach exceeds ICC for other approaches

| | Mean | 90%<br>CI<br>Low | 90%<br>CI<br>High | Interval<br>Width | $P(ICC_{mixture} > ICC)$ |
| --- | --- | --- | --- | --- | --- |
| **Mixture** | 0.754 | 0.668 | 0.802 | 0.133 | n/a |
| **Expanded** | 0.730 | 0.658 | 0.775 | 0.117 | 0.744 |
| **Up** | 0.721 | 0.563 | 0.777 | 0.214 | 0.739 |
| **Down** | 0.704 | 0.567 | 0.767 | 0.199 | 0.823 |
| **Random** | 0.682 | 0.452 | 0.755 | 0.303 | 0.744 |

### Intra-Rater Reliability

The intra-rater ICC for fibrosis staging for the *Mixture* model was around 0.791 (CI: 0.757-0.826). The probability that the ICC estimate for the *Mixture* model surpassed other *ad hoc* ICC estimates was as high as 86%, while ICC estimates given by the *Expanded* model were more similar to the *ad hoc* procedures. For instance, the random staging ICC estimate was found to be approximately 5% lower than using the *Mixture* approach (Table 3). The posterior intervals of ICCs for the *Bridge Category Model* approaches (Mixture, Expanded) were significantly lower than the *ad hoc* procedures, indicating greater precision ^39^.

**Table 3:** Intra-Rater ICC estimates for all statistical approaches and probability of how much ICC for *Mixture* approach exceeds ICC for other approaches

| | Mean | 90%<br>CI<br>Low | 90%<br>CI<br>High | Interval<br>Width | $P(ICC_{mixture} > ICC)$ |
| --- | --- | --- | --- | --- | --- |
| **Mixture** | 0.791 | 0.757 | 0.826 | 0.069 | n/a |
| **Expanded** | 0.763 | 0.730 | 0.799 | 0.069 | 0.830 |
| **Up** | 0.767 | 0.727 | 0.832 | 0.106 | 0.790 |
| **Down** | 0.769 | 0.731 | 0.816 | 0.086 | 0.766 |
| **Random** | 0.752 | 0.710 | 0.826 | 0.117 | 0.860 |

### Bridge Category Analysis

When pathologists assign bridge ratings under the assumptions of the *Mixture* Bridge Category Model, they originally have a definitive stage in mind for prognosis. To hedge against the possibility of a higher or lower stage being true, they may blur the rating into two adjacent categories, generating the bridge rating. Given the assignment of a bridge rating, a pathologist may feel the lower of two assigned adjacent categories is the truth *p* proportion of assignments and the higher of the two assigned adjacent categories *1-p* proportion of assignments, which demonstrates a decision-making processes during staging. Studying decision making processes for specific pathologists can inform optimal staging practices.

Pathologists demonstrated different propensities for believing the lower or higher stages to be true with regards to the bridge category assignments under the *Mixture* Bridge Category Model (Figure 1; Table 4). For test 1, pathologist 1 expressed that he/she felt that the true stage was the lower of the two adjacent stages 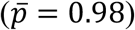, definitively greater than the non-preferential 0.5 proportion (1-pd=0.001), as opposed to pathologists 2 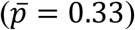, who leaned towards expressing the higher of the two adjacent as the true stage. For test 2, pathologist 1 did not assign any bridge ratings, such that the posterior interval covered all values from 0-1 to indicate lack of information regarding bridge category preference. Meanwhile, pathologist 2 demonstrated the preference for the lower of two stages 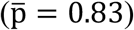, a result definitively larger than 0.5 (1-pd=0.02). Both pathologists 2 and 4 exhibited different bridge category preferences between the first and second test, yet pathologists 3 and 4 did not demonstrate preferences in any particular direction throughout the tests.

**Table 4:** Breakdown of *mixture* proportion *p* by pathologist and measurement of how proportions compare to lack of bridge rating preference (*p*=0.5) and changes in preference versus the previous timepoint

| | | Mean | 90% CI Low | 90% CI High | Posterior Interval | 1-p-direction, versus $p=0.5$ | 1-p-direction, versus Test 1 |
| --- | --- | --- | --- | --- | --- | --- | --- |
| **Test 1** | Pathologist 1 | 0.98 | 0.89 | 1.00 | 0.11 | 0.001 | n/a |
|  | Pathologist 2 | 0.33 | 0.00 | 0.63 | 0.63 | 0.191 | n/a |
|  | Pathologist 3 | 0.62 | 0.36 | 0.85 | 0.49 | 0.218 | n/a |
|  | Pathologist 4 | 0.42 | 0.20 | 0.64 | 0.44 | 0.280 | n/a |
| **Test 2** | Pathologist 1 | 0.48 | 0.00 | 1.00 | 1.00 | 0.476 | 0.188 |
|  | Pathologist 2 | 0.83 | 0.56 | 1.00 | 0.44 | 0.020 | 0.004 |
|  | Pathologist 3 | 0.46 | 0.15 | 0.72 | 0.57 | 0.417 | 0.155 |
|  | Pathologist 4 | 0.65 | 0.44 | 0.89 | 0.46 | 0.132 | 0.090 |

**Figure 1:**
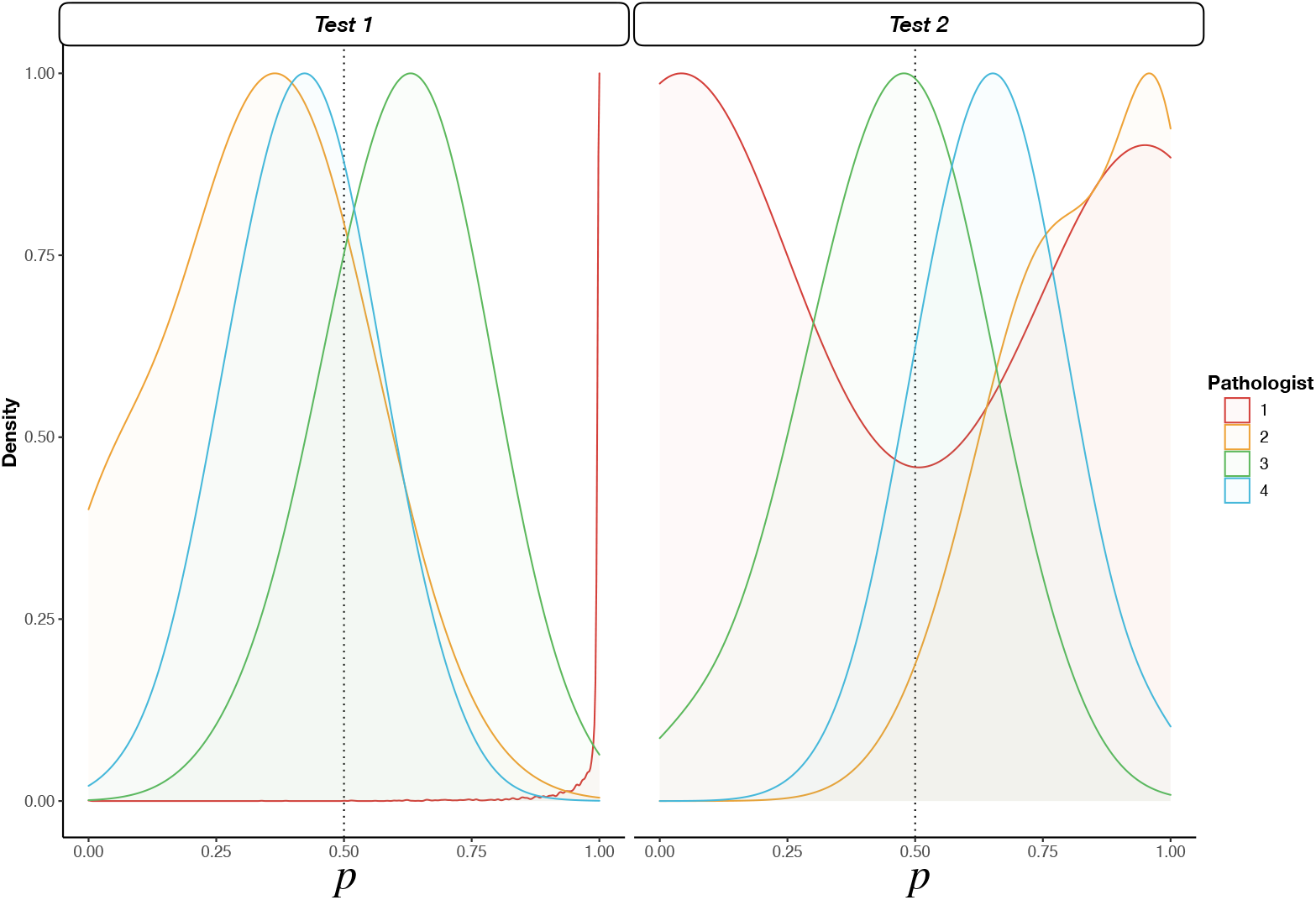
Density plot of 1000 posterior draws for *mixture* proportion *p* for Tests 1 and 2 by Pathologist.

## Discussion

The results from our study indicate that failing to take into account bridge ratings could potentially reduce inter-rater ICC estimates for NASH Fibrosis by up to 10% and reduce the intra-rater ICC by up to 5%. Given that ICC scores assigned to the *Mixture* model trended upwards versus other approaches, while ICC scores assigned to the random approach trended downwards, this suggests that the former technique’s capacity to account for latent heterogeneity, while random assignment of bridge ratings exacerbates heterogeneity, thereby driving reliability downwards. The magnitude of the effects is not surprising, since bridge categories reflect movements of stages over small scales and should not impact variation as much on a 5-point scale.

Additionally, utilizing *Bridge Category Models* yielded significantly higher precision for the ICC estimates versus *ad hoc* procedures. These significant precision gains translate to greater confidence about the scale’s reliability when communicating the study findings, as the statistical models may provide a better fit to the data since they better represent the data generating mechanism.

Additionally, we demonstrated that pathologists exhibit different preferences when assigning bridge ratings and sometimes contradict themselves in their assignment across tests. Estimates of a rater/test specific proportion *p* provides nuanced information on staging practices with respect to bridge ratings by allowing comparisons between pathologists over time to be made, potentially informing future staging practices. For instance, pathologist 1 clearly demonstrated the propensity for assigning two higher stages for bridge ratings, which indicates that they felt biopsy features are suggestive of the lower stage but hedges against the possibility of a more severe outcome.

NASH is becoming an increasingly widespread condition around the world and necessitates the adoption of assessment methods that have greater fidelity to the underlying data generating mechanisms to allow an honest and transparent assessment of clinical drug trials and artificial intelligence (AI) technologies^9,20,40–42^. While the magnitude of the study findings appears to be dwarfed by the overall clinical impact of lower measurement reliability, being able to ensure that our measurement scales are properly calibrated to these issues is immensely important; small percentage differences multiplied across a large number of cases can heavily impact care. These results, irrespective of the modeling approach, offer further confirmation regarding the generalizability of the NASH CRN scale in that significantly high reliability was not achievable, either between or within pathologists. These findings are consistent with a crop of new studies which have disputed the generalizability of the original NASH scale establishment ^9–16^.

While our results further support the lack of generalization of the NASH CRN scale, our study is limited in that stage was assessed via inspection of Whole Slide Images, of which extensive research does not exist to verify the concordance between assessment via microscope or computer ^43–48^. The pathologists also had an appropriate washout period during the first and second time they viewed the specimen, however, this does not preclude the possibility that some features of the initial biopsies may have been remembered from the previous inspection, which may have influenced some of the rater-specific effects for *p*. We did not account for heteroskedasticity or category-specific effects, where the variation in staging or *p* may change based on how close it is to zero or one as a function of the fixed and random effects ^36^. Nor did we estimate bridge category effects on a case-by-case basis, though we have provided such a model formulation in the Appendix (section “Alternative Mixture Model Specification to Account for Biopsy-Specific Bridge Category Effects”). We note here that our models only estimate reliability and variation but not the treatment effect, from which prior studies have demonstrated that failing to account for bridge categories can lead to great bias in the study conclusions. However, we have not yet tested for correlates with fibrosis stage (e.g., effect of drug on fibrosis regression, serological correlates, assessment of AI technologies) in the hierarchical setting, which presents a future area of inquiry.

## Conclusion

In this work, we applied hierarchical Bayesian *Bridge Category Models* to resolve bridge ratings for the assessment of the generalizability of the NASH CRN scale. Estimates of the reliability of the scale trended upwards when using the *Mixture* model versus previous *ad hoc* approaches, while reliability trended downwards for random assignment of bridge rating. Additionally, *Bridge Category Models* may allow researchers to communicate reliability findings for fibrosis staging with greater precision. Consequently, using *Bridge Category Models* over the *ad hoc* approaches increase the confidence that bridge ratings would not negatively impact or bias study conclusions by overcoming potential deficiencies associated with *ad hoc* procedures. In this study, we demonstrated that their hierarchical incarnation allows for the assessment of how pathologists differed in their treatment of bridge categories to potentially inform future staging practices. Using our *Bridge Category Models*, we provide further confirmation of the lack of generalizability of the NASH CRN scale. We expect these hierarchical Bayesian models to be more applicable in the estimation of a treatment effect (e.g. regression of Fibrosis stage), where small changes multiplied across large cohorts could improve the power of clinical research studies on NASH drugs. As such, we plan to continue to assess how these reliability and uncertainty issues impact clinical trial design and evaluation as well as the development and validation of AI technologies to assess in the evaluation and staging of NASH.

## Appendix

### Additional Description of Bayesian Modeling Procedures

Bayesian model fitting procedures estimate posterior distributions for each covariate effect and combinations thereof by coherently updating the information expressed in the prior, representing prior knowledge about the effects (often unassuming about the likelihood of one value versus another), and the experimental information in the likelihood function, which expresses the likelihood of observing the data given the value of the effects or hypotheses involving them. Bayesian methods estimate the complete posterior distribution of the effect as the primary analytic summary, whereas frequentist approaches often produce approximate inferences based on the sampling distribution of the estimator used to estimate the effects. In more complication situations, frequentist procedures may be biased or fail to converge. The prior in Bayesian statistics may reduce the expected error of a Bayesian procedure by shrinking the effect estimate towards a more conservative value (resembling penalization procedures in frequentist statistics). The posterior from one analysis may become the prior for a subsequent analysis, portraying the natural updating or learning formula that is characteristic of Bayesian analysis. The fundamental element of the Bayesian approach is the expression:

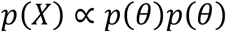

where *p(θ)* is the prior, *p(θ)* the likelihood, and *p(X)* the posterior or updated beliefs. Markov Chain Monte Carlo (MCMC) procedures, or other popular approaches (e.g., Stochastic Variational Inference, Quadratic Approximation, Hamiltonian Monte Carlo / HMC) are used to sample parameter values from the posterior. MCMC procedures iteratively sample the posterior one parameter at a time, conditional on the other parameters (i.e. the Markov Chain) In contrast, HMC approaches may sample the joint posterior via auto-differentiation procedures ^21,23,30,31^. A cumulative link model, the ordinal regression model used to calculate the ICC, may be formulated in terms of the prior and likelihood as follows:

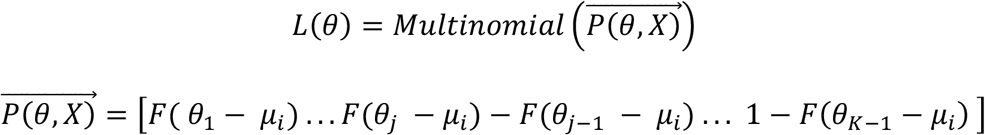

with *F*() the cumulative distribution function of the standard normal distribution,

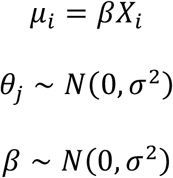

Where additional hyperpriors may be specified for *σ*^*2*^. Note that this model, which is the basis of our prior work ^17^, fails to account for nested observations.

### Complete Bayesian Specification of Hierarchical Mixture Bridge Category Model

Hierarchical modeling procedures allow for flexible modeling of nested observations, where statistical dependencies exist between observations. Here, multiple measurements are rendered per case/biopsy across multiple raters, where each rater examines the biopsy more than once. This dependency structure places repeat stage assignments closer to each other if they came from the same rater/case. If ignored, the effect estimate for an ordinal regression model may be confounded/biased by batch and spurious findings may result. If averaged across other observations in the batch, variation is removed from the data, leading to a risk of overstating the model fit and significance of the association. To resolve these issues, random intercepts per cluster of observations are added to capture baseline differences in measurements, while random slopes may capture time-dependent differences in measurements within clusters, as an example.

Random intercepts over the clusters are assumed to be drawn from a population of intercepts with some variance, 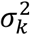, where *k* could be the factor being measured across (e.g., *k* represents patients), while observations within a patient vary according to an amount of variation given by 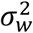. For validation of measurement scales, often it is important to understand the variation between 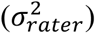 and within raters 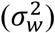, a measure of the scale’s reliability and a proxy for measurement error.

Derivation of ordinal regression models of the cumulative link and Bridge Category models have been described in a previous work ^17^. A hierarchical extension of these procedures enables estimation of the aforementioned variance components. The likelihood function for the entire sample of observations is given by:

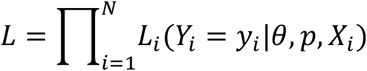

with the contribution from the ith observation given by:

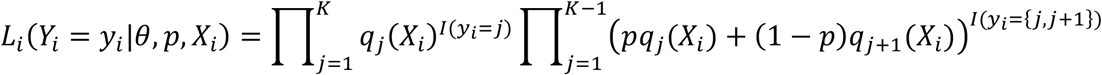

where:

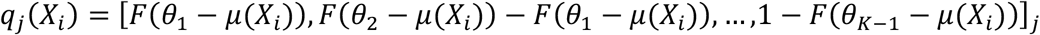

with *F*() the cumulative distribution function of the standard normal distribution,

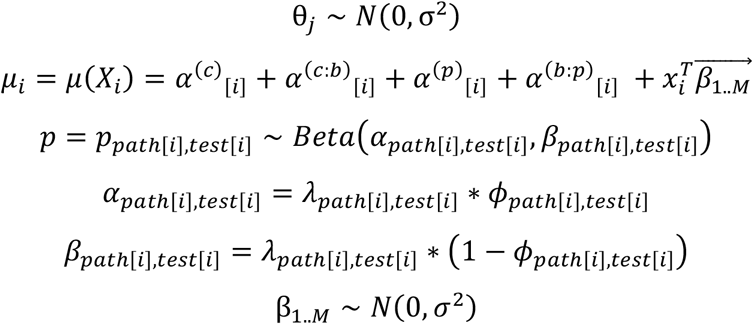

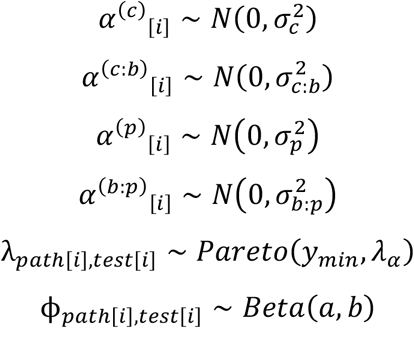

First, it is determined whether a bridge rating, *I*(*y_i_* = {*j*,*j* + 1}), has been assigned to the observation. The rater/test specific mixture probability, *p*_*path[i]*,*test[i]*_, is selected after indexing by pathologist (*path*) and test. The likelihood weighs information across all categories as determined by the presence of a bridge rating and the drawn mixture parameter. If there is no bridge rating, the likelihood is then set to attribute fully the contribution from category *j*. Finally, the likelihood weights, are multiplied elementwise by the multinomial probabilities, 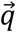, across all possible response categories, which are added together to yield the likelihood *L*_*i*_*(Y*|*θ)*.

The components of variation in staging attributed to different patients/cases is 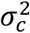, and that of biopsies nested within patients is 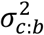 The total variance attributed to the biopsies are given by:

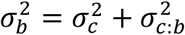

Furthermore, given selection of a particular biopsy, the variance in assigned stage attributed to pathologists are given by 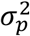and the variance attributed to the cross-classified interaction between pathologist and biopsy is 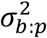.

The *Mixture* parameter *p*is estimated on a pathologist and test basis, given by parameter matrix 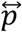, comprised of four pathologists (columns), each completing two tests (rows). Mean and scale hyperparameters *ϕ* and *λ* are reparametrized to form a beta prior for each pathologist/test-specific *α* and *β*. Since the pathologist/test specific *mixture* parameters are separately estimated once per HMC iteration, which allows separate priors for *ϕ* and *λ* that are more flexible priors/hyperpriors for *p*. Furthermore, hyperpriors may be set over the variance components.

### Alternative Mixture Model Specification to Account for Biopsy-Specific Bridge Category Effects

As an alternative model specification, we can model pathologists and their interaction with test number (i.e., did bridge rating propensity for down/upstaging *p*vary by pathologist, over number of repeat tests?) by estimating the relationship between covariates 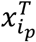 and conditional mean *μ*_*pi*,_ which maps to *p*, rather than estimating separate intercept terms for the different effects:

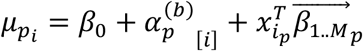

The conditional mean increases monotonically with *p*, such that higher values of a covariate for positive parameter values will lead to higher reports of *p*. Repeat measurements over biopsies are accounted for via adoption of random intercepts 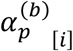. The covariates are fixed effects by rater/test number:

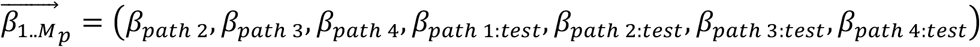

For the *Mixture* bridge rating models, these variance components may also be estimated to account for the impact of uncertainty in them on inferences for *p*, the propensity for up/down-staging. In addition, fixed effects components 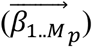, a new set of bridge-rating specific covariates, 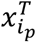, may also be estimated. In this case, we treat pathologist as a fixed effect for the estimation of *p*. As such, we present the following prior and likelihood specifications for an alternative hierarchical extension to the Bridge Category Models, in the case of the *Mixture* approach:

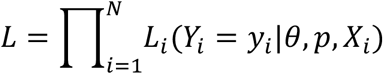

with the contribution from the ith observation given by:

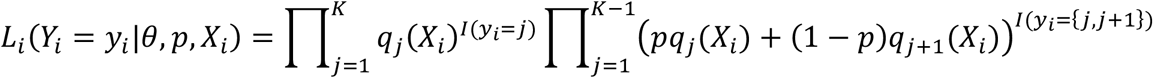

where:

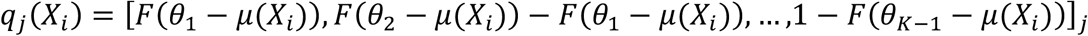

with *F()*the cumulative distribution function of the standard normal distribution,

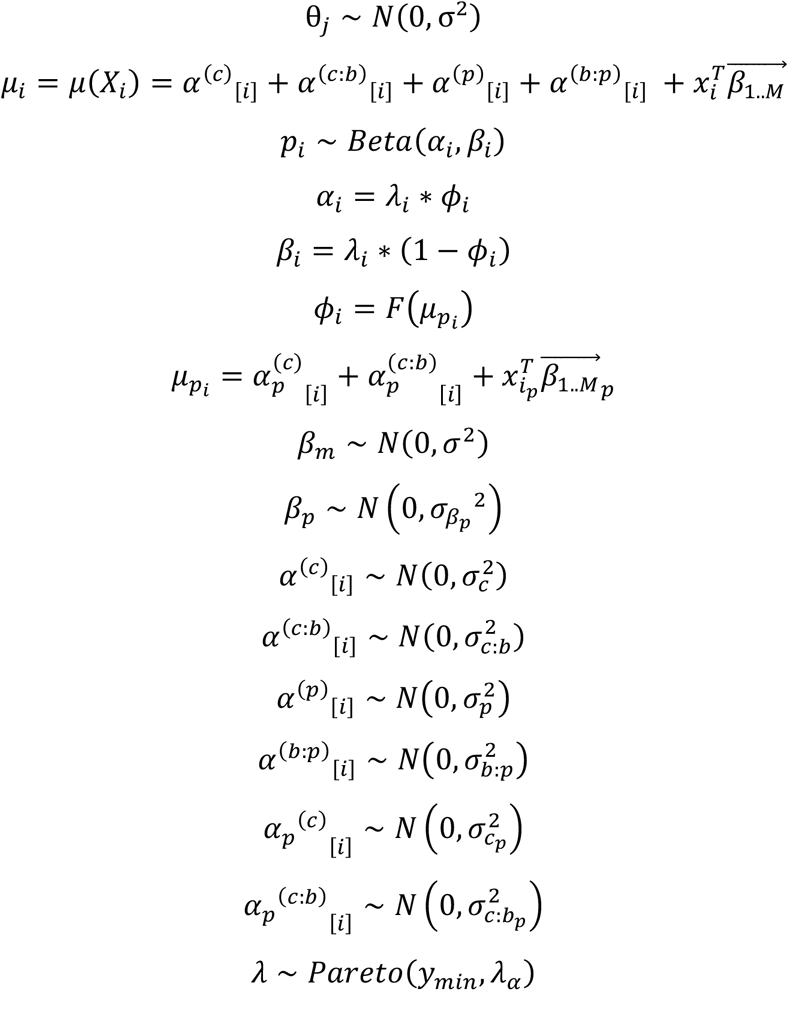

This statistical model adds random intercepts for cases *c*, nested biopsies *c:b* and fixed effects for pathologists *p*, not to be confused with the mixture parameter *p*, where appropriate as mentioned in the main text. Additional random intercepts are used to estimate variance components for the estimation of the mixture parameter *p*, while either fixed or random effects can be used to compare factors relating to the estimation of *p*. The cumulative distribution function *F* maps the conditional means to a value between 0 and 1, which is helpful for generating values of *p*from a beta distribution by incorporating a valid mean parameter *ϕ*for such a distribution.

### Inference Targets

For the assessment of inter and intra-rater ICC, the main target of inference is the vector of variance components:

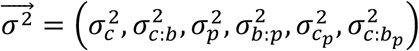

To understand how some of the covariates *X* relate to stage assignments / bridge category up/down-staging, the relevant quantities are:

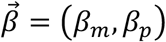

To understand how pathologists differ in their stage assignments, pathologist identifiers may be modeled as either a fixed or random effect; in this work they were modeled as a random effect to estimate pathologist level variance. For our work, we also set all of the staging covariates to 1 (*X= 1*) to capture the population level intercept, which does not impact estimation of the variance components around the population intercept.

The pathologist-specific *mixture* parameters were estimated for each test number (indicator matrix) as bridge rating specific terms for the *Mixture* model:

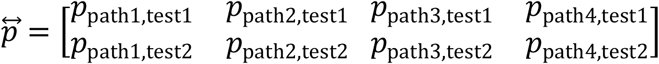

### Description of Priors

We used the following priors for our hierarchical Bayesian models:

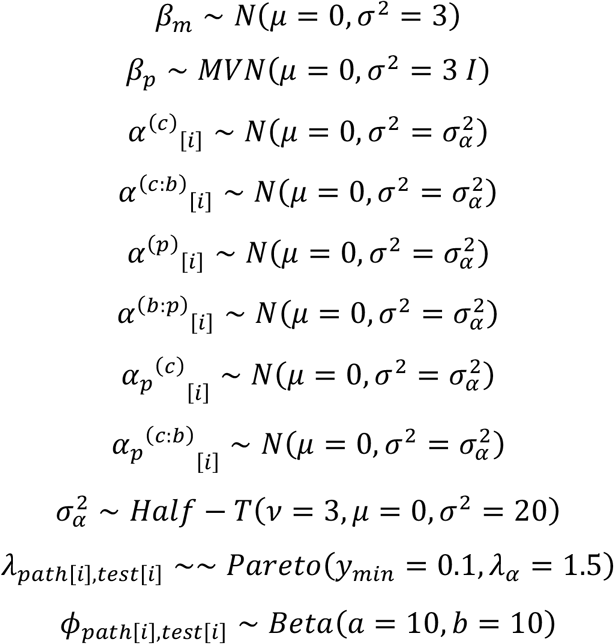

A conservative prior was used for *β*, while a weakly informative prior has been specified for the random intercepts. The hyperprior over 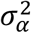 actually represents the individual priors for each of the six sets of random intercepts being generated:

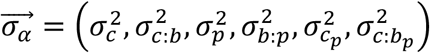

## Notes

### Competing Interest Statement

The authors have declared no competing interest.

